# Characterizing mechanics of membrane tether from relaxation curve obtained by optical tweezers

**DOI:** 10.1101/2020.09.06.285353

**Authors:** Xuanling Li, Xiaoyu Song, Yinmei Li, Ming Li, Haowei Wang

## Abstract

Optical tweezers is a powerful tool in the study of membrane tension. Comparing to pulling out an entire membrane tether at one time, the step-like method is more efficient because multiple relaxation curves can be obtained from one membrane tether. However, there is few proper models that describe relaxation curves to characterize mechanical properties of cell membrane. Here we established a model to describe the relaxation curve of HeLa cells based on the relationship between membrane tether diameter and tensions. We obtained effective viscosities and static tensions by fitting relaxation curves to our model. We noticed the delicate structure of relaxation curves contains information of cell skeleton changes and protein diffusion. Our study paved a novel pathway to characterize the dynamics and mechanics of cell membrane.

---

Cell membrane is an active and dynamic structure. The tension and curvature of cell membrane, as well as the protein density on cell membrane, are the basis of physiological functions, such as membrane transport, cytoplasmic division, infection and immune response^1,2^ The cytoskeleton, membrane protein, and membrane tension regulate many significant biological processes^3,4^, which are very important for cells to respond to external mechanical stimulation^5^. For example, the attachment/disassembly at the front/rear of cells regulated by membrane tension constitutes the basis for cell migration^6,7^. Cell secretion and endocytosis also involve cell membrane tension change^8^. The study of cell tension and the interaction between membrane proteins and cytoskeleton has become a hotspot for cell surface mechanics^9^.

Stretching membrane tethers from cell membranes had been recognized as standard method for cell surface mechanics study^3,4,9^. There are two direct methods to produce membrane tethers: use a flowing stream to push cells that pre-attached to the underlying surface^10^, or stretch membrane tethers with a modified AFM probe^11,12^. In addition, membrane tethers were also extracted from cells with modified beads manipulated by micropipette^13,14^, optical tweezers^4^, and magnetic tweezers^15^. Among these techniques, optical tweezers has been widely applied to study membrane tethers of neurons^16^ and neutrophils^7^, and to find membrane reservoir inside mouse fibroblasts^17^.

Membrane tether elongation requires a continuous stretching force to overcome both membrane tension and the resistance caused by the viscous phospholipids flow from the cell membrane to the membrane tether. Most of studies on membrane tether focus on the minimum force required to maintain a tether *f_0_* (static force) and the coefficient of effective viscosity *η_eff_*. It is well accepted that *f_0_* is the measure of membrane tension and *η_eff_* corresponds to the friction between lipid bilayer and membrane skeleton. However, the only experimental observations are curves of force and tether length *vs*. time. Therefore, researchers are in great needs of physical models to find out *f_0_* and *η_eff_* from experimental data^4,12,18–20^.

At present, commonly used methods are stretching out membrane tethers at different speeds, then fitting tensions and velocities into a straight line, to calculate the overall tension and effective viscosity coefficients^14,21–23^. However, there are three drawbacks in this strategy:

1. The model requires pulling out entire membrane tethers at one time at different speeds. Fitting stretching force and speed might be impracticable, if the upper limit of stretching speed is small.
2. The model requires membrane tethers several microns or longer in order to obtain stable data^24^. However, the pulling force increases sharply upon membrane reservoir depletion within couple of micro-meters^25^. Furthermore, long membrane length and stretch time also mean high possibilities of broken tether during experiments, resulting low efficiency. The number of stretches increases in the case of collecting stretching forces at different speeds. It further reduces the experimental efficiency.
3. Current methods only use overall pulling force and speed, and ignore stretching curve details. However, these details may contain abundant information, such as how the cytoskeleton breaks, and how membrane proteins diffuse from cell membrane to membrane tether during tether elongation.

Obviously, stretching a membrane tether in a step-like manner repeatedly is more practicable than regular strategy. Previous studies noticed the tension decreased approximately exponentially after active stretching^26,27^. However, this strategy has not been widely used because of the lack of suitable models. In this paper, we established a physical model to analyze relaxation curves obtained during step-like stretching, based on well accepted relationship between force (*f*) and elongation speed (*V_t_*)^23,28–31^. Comparing to the former method, our model has significant improvements as following: 1. One single experiment can provide dozens of relaxation curves. Each curve provides a pair of *f_0_* and *η_eff_*. Both greatly improves the efficiency. 2. Improved efficiency enable people to study statistical correlation of *f_0_*, *η_eff_* and tether length. 3. The step-like method has no requirement on tether stretching speeds. That expands using cells with tethers that are difficult to stretch out. 4. The new model is capable of analyzing dynamic information of protein diffusion by studying relaxation curves. In summary, our physical model not only greatly improves the efficiency of measuring membrane filaments, but also provides a novel approach to the studies of mechanical properties and dynamics of cell membrane.

## Results

The optical tweezers remained stationary during experiment. We first used optical tweezers to capture a 2 μm PS bead and hold it 2 μm above the bottom of the chamber. Then, a HeLa cell attached to the bottom of the chamber was moved towards the captured bead by piezoelectric stage. The bead and the cell contact for 5-10 s. Then the cell was moved 0.5-2 μm away from the bead at the speed of 1 μm/s to observe whether membrane tether was formed. Once the membrane tether was confirmed, the tether was elongated in a step-like manner: The cell first moved away from the bead 0.5-1μm in 0.5 second as the tension increased dramatically to its maximum value (Fig. 1A). Once the cell stopped moving, the tension began to decrease and the bead was pulled back by optical tweezers (Fig. 1B) until equilibrium (Fig. 1C). The tension *vs*. time was recorded for analysis (Fig. 1).

**Figure 1.**
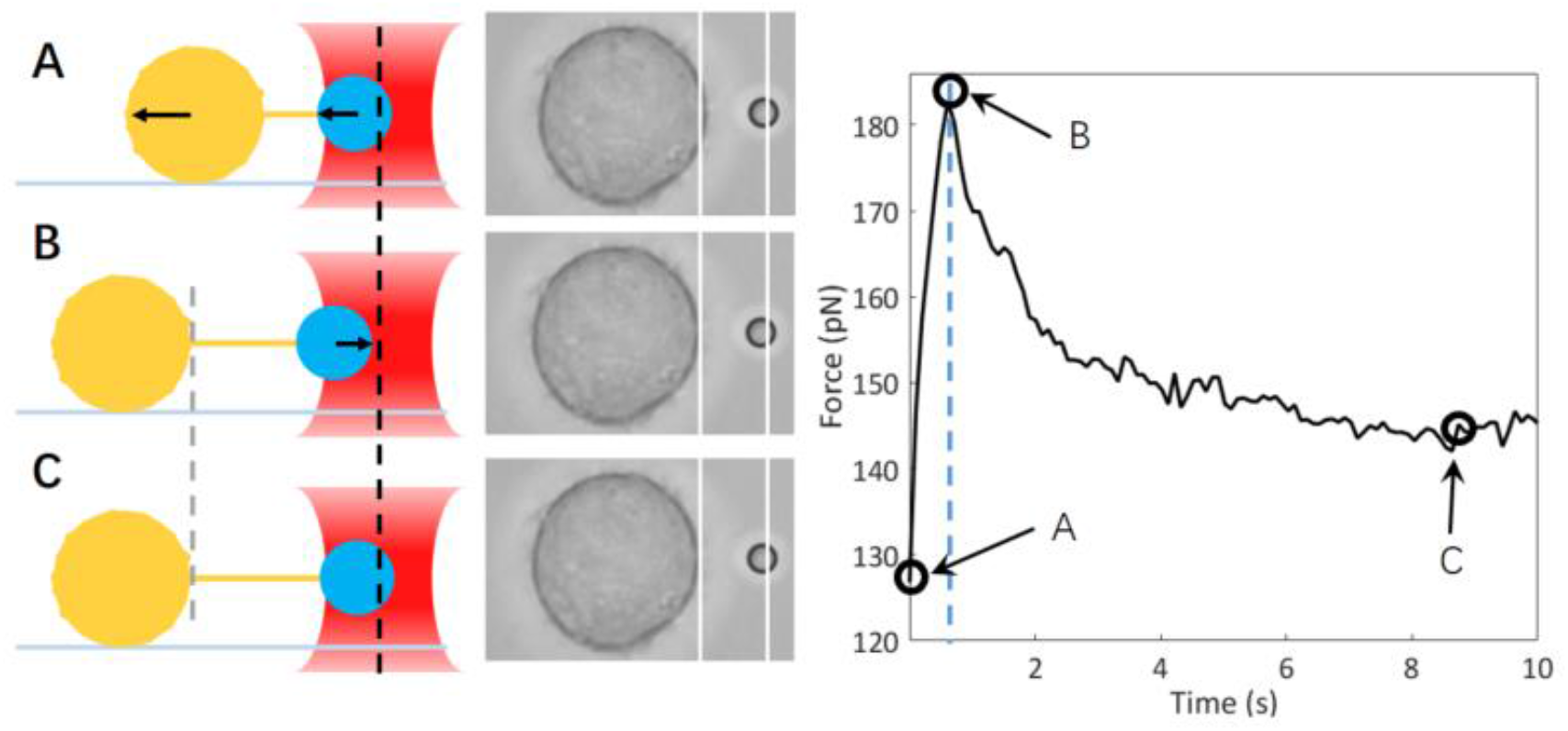
Illustration of stretching membrane tether of a HeLa cell with optical tweezers, and the tension relaxation curve. The left figure are schematic drawings of the experiment, in which big circles represents HeLa cell, and the small circles represents PS bead captured by optical tweezers; the line between the two circles represents the membrane tether, and the arrows indicate motion directions. (A) The cell moves to the left and membrane tether elongates. The PS bead is pulled left due to increased tension. (B) Once the cell stops moving, the bead is pulled backward towards the center of the optical tweezers. (C) The bead stops moving until the tension decreases to f0. In the middle is the screenshot of each situation in experiment. The right figure shows the relaxation curve of the tether tension with time during the stretching process, in which the black circles correspond to the time from A to C.

We obtained 109 relaxation curves from HeLa cells in experiment (Fig. 2A). The final tensions were in between 100-150 pN (Fig. 2). We choose maximum tension *f_s_*, effective viscosity coefficient *η_eff_* and static tension *f_0_* as fitting parameters (Material and Methods, equation (10)). Most of relaxation curves in the experiment are consistent with our model (Fig. 3A). However, there are obvious turning points in some of the curves. We divided those curves into sections before and after the turning point (Fig, 3B) and found that the *η_eff_* (51.72 vs. 6.52 pN•s/μm) and *f_0_* (70.0 vs. 52.4 pN) of the two sections are significantly different. For curves contain one or more turning points, at least one of *η_eff_* and *f_0_* change significantly after the turning point, which indicates abundant dynamic changes in the process of membrane tether elongation (Figure S1, Table S1).

**Figure 2.**
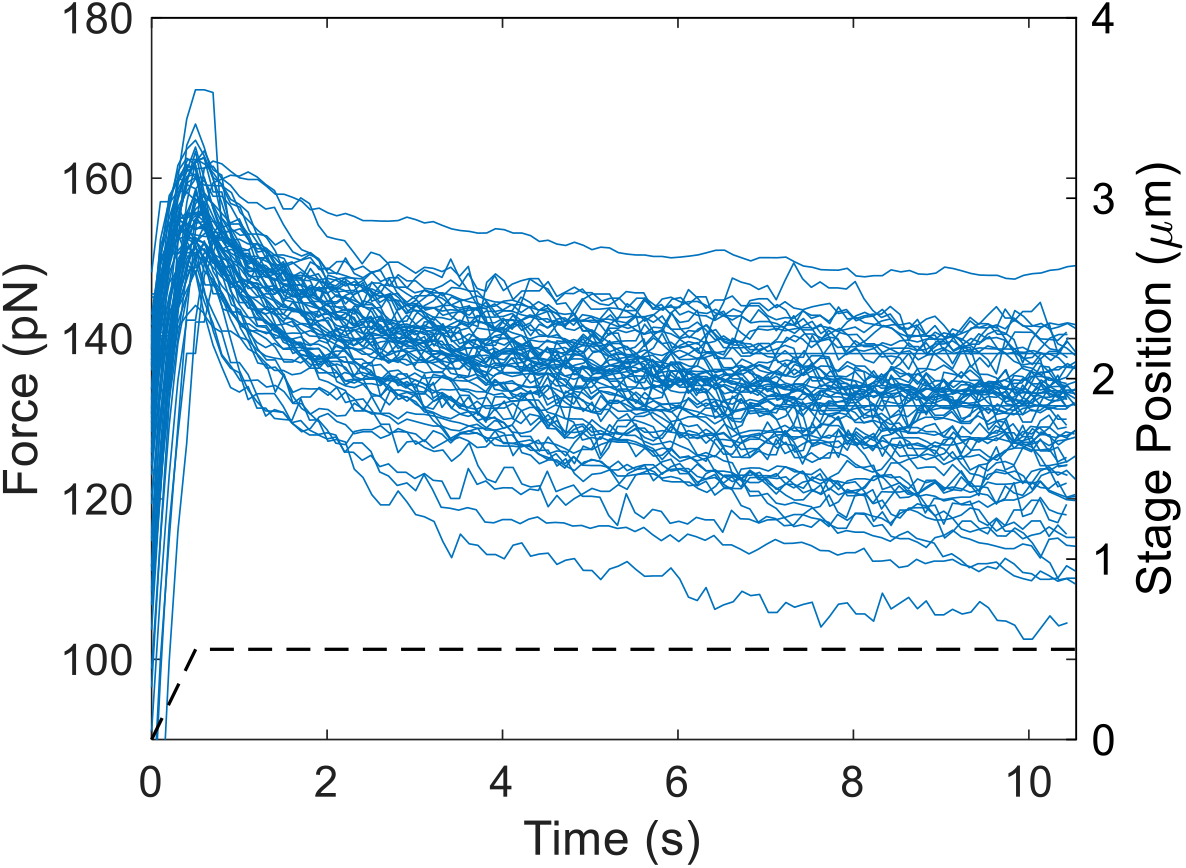
Relaxation curves of membrane tethers stretched from HeLa cells. The left coordinate is the tension. The right coordinate and the dash line represent the positon of PZT stage.

**Figure 3.**
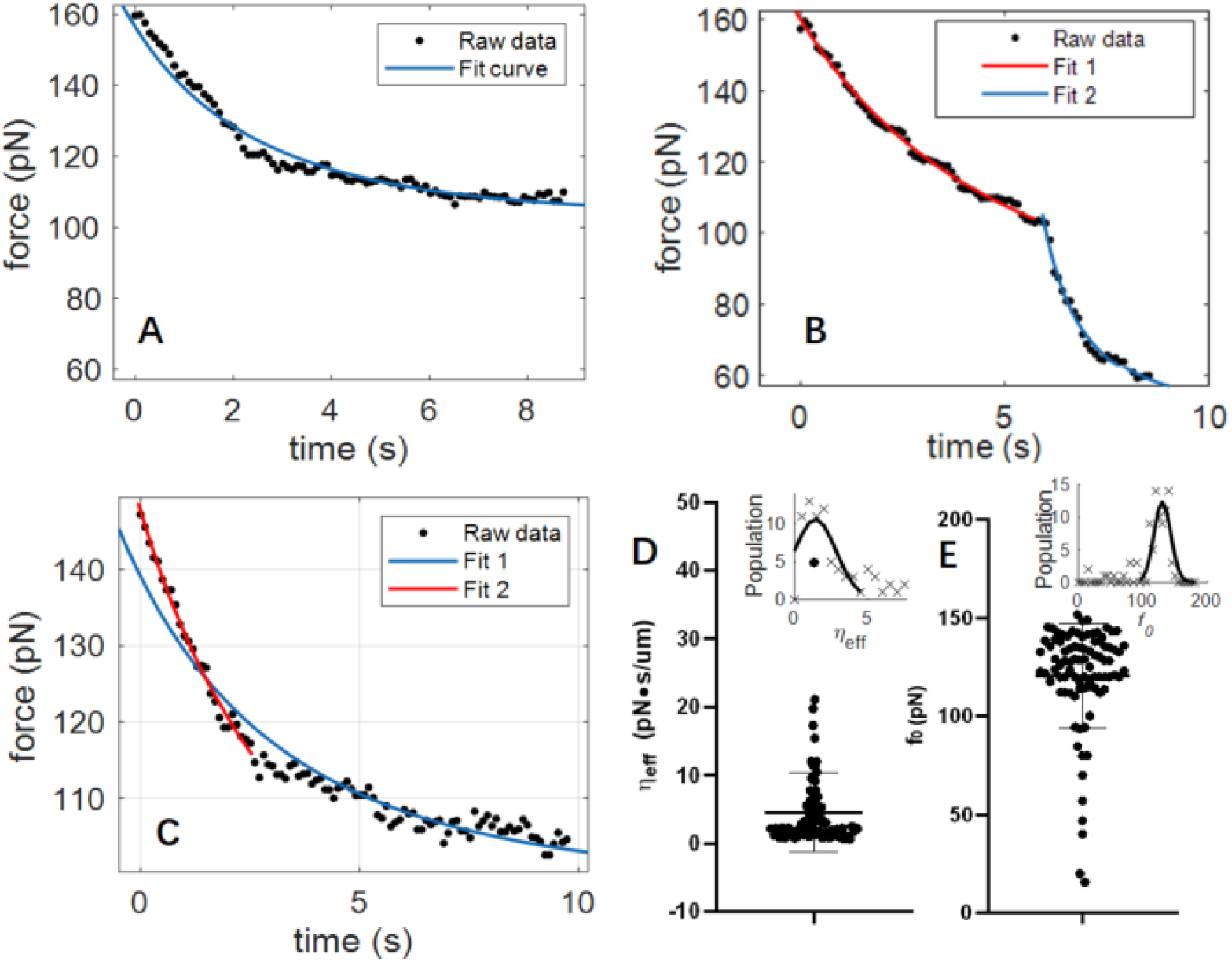
Characteristics of HeLa cell membrane tension. (A) a set of relaxation data and fitting curve. (B) A relaxation curve carrying a turning point in the middle. The left and right section of curve were fitted with equation 10 respectively. (C) When the tether length exceeds 10 μm, only the first two seconds data consistent with the model in some curves. The blue solid curve is the result of fitting all data. The red line is the fitting result with first two seconds data only. (D&E) The distribution of effective viscosity *η_eff_* and static force *f_0_*. The inset curves are Gaussian fitting of *η_eff_* and static force *f_0_* distribution.

When tether length exceeds 10 μm, about 50% of the relaxation curves deviated from our model significantly. However, curve of the rapid drop section before the turning point, still consistent with our model (Fig. 3C). After the turning point, the tension decreases slowly, following a linear-like trend. In order to eliminate the deviation after the turning point, we intercepted the first two seconds data of the curve and fitted out the *η_eff_* and *f_0_* (Fig. 3D&E). We found lots of *η_eff_* concentrated at 1.5 pN•s/μm, forming a plateau; *f_0_* mainly distributed between 110-150 pN (Fig. 3D&E inset, Table 1).

**Table 1:** Fitting results of 0-2s relaxation data

| $\eta_{eff}$ ($\text{pN}\cdot\text{s}/\mu\text{m}$) | | $f_s$ (pN) | | $f_0$ (pN) | |
| --- | --- | --- | --- | --- | --- |
| Mean | Standard deviation | Mean | Standard deviation | Mean | Standard deviation |
| **4.56** | 5.80 | 153.98 | 12.55 | 120.68 | 26.56 |

Then we looked at the correlation of distributions between *η_eff_, f_0_* and tether length. We found the population of larger *η_eff_* decreased as the tether length increase (Fig. 4). When the length of the tether is more than 10 μm, *η_eff_* concentrated near the minimum value with few large values. In contrast, the distribution of *η_eff_* is dispersed when tether length is less than 10 μm (Fig. 4 inset). In addition, when the tether length is longer, *f_0_* tends to concentrate between 110-115pN, and the population of small *f_0_* is rare (Fig. 4B). However, there is no significant correlation between *f_0_* and *η_eff_* (Fig. 4C), indicating that the two parameters are not significantly correlated.

**Figure 4.**
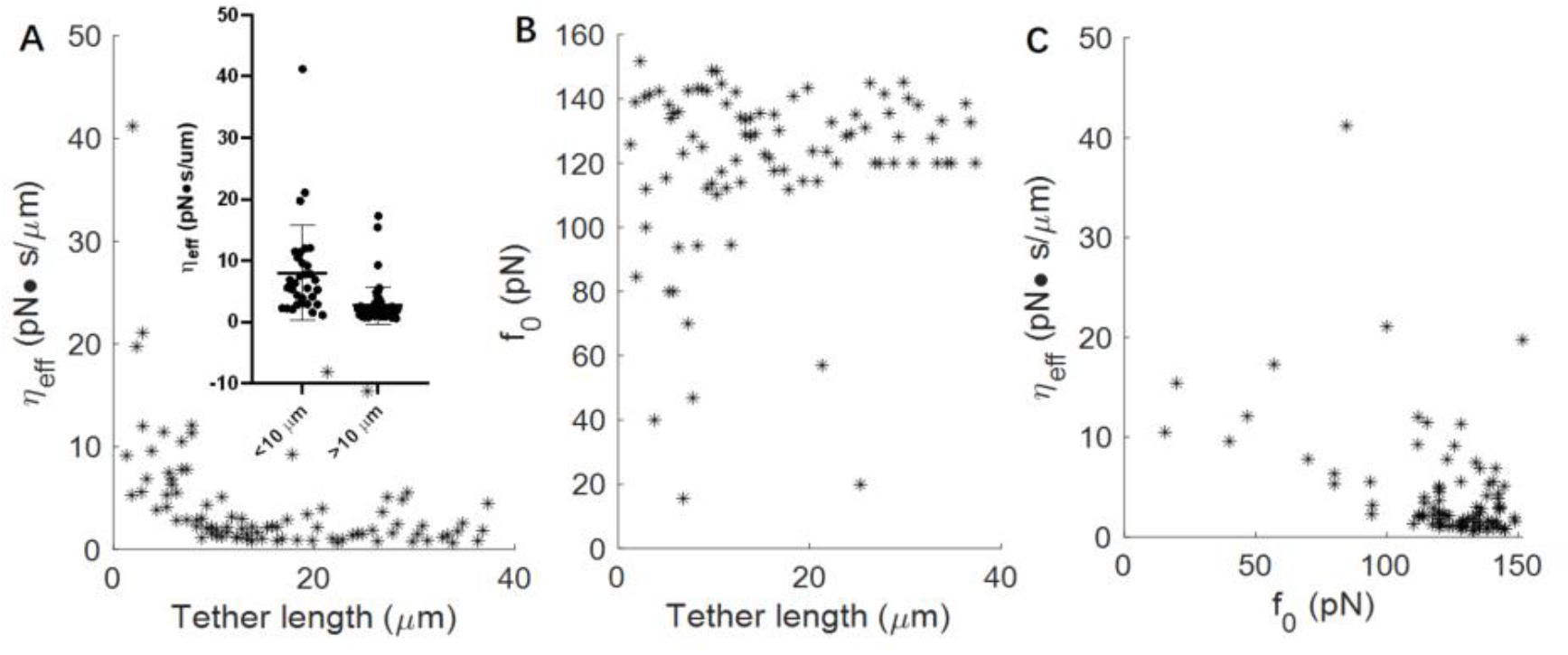
Statistical relationship between *η_eff_, f_0_* and tether length observed from HeLa cell membrane tether. (A) Correlation distribution between *η_eff_* and tether length; The inset illustration shows the distribution of *η_eff_* when the tether length is shorter and longer than 10 μm. (B) Correlation distribution between *f_0_* and tether length. (C) Correlation distribution between *η_eff_* and static tension *f_0_*.

## Discussion

We obtained over 100 relaxation curves from HeLa cells with step-like method. The membrane tether only extended 0.5-1 μm every time. Each measurement took about only 10 seconds and rarely fails. Compared with the widely used method (continuous stretching and fit the force *vs*. speed plot with a linear model), the experimental efficiency has been greatly improved and can be used to study cells that are difficult to pull out long tethers.

The key of step-like method is to find a proper physical model that can characterize mechanical properties from relaxation curves. In 2015, Pullarkat *et.al*. represented a two-component model to describe relaxation curves obtained from Chicken dorsal root ganglia (DRG) neurons^16^. They hypothesized that the membrane contains at least two components with very different amount, and the neck region at the base of the tether acted as a potential barrier for the minor component. Therefore, if one assumed most of minor components did not transport to the tether during stretching, the diffusion of this minor components could be used to explain the relaxation behavior right after stretching.

Although Pullarkat’s model can be used to fit experimental data, there is no direct evidence that the minor component is involved in the tension decline process in relaxation curves. Furthermore, Pullarkat’s model hypothesized there was a membrane composition gradient from the axon to the tether, but they did not give a convincible mechanism of how such gradient came into being. In fact, it is hard to image the minor component does not transport from membrane to tethers during stretching. Most importantly, in 2013, Skykes *et. al*. reported similar relaxation behavior with several seconds timescales from tethers made of pure lipids^32^. Therefore, it is reasonable to suspect the diffusion of minor components does not dominate the relaxation behavior.

The reason to introduce effects of multiple components is that published timescales of relaxation curves of single component tether does not exceed 10^-2^ seconds, which can hardly explain the tension decline in several seconds^16^. According to our model, the tension could decrease to *f_0_* under a reasonable *η_eff_* within couple of seconds, even for one component membrane tethers. Therefore, we believe the extension of tether radius dominated the relaxation behavior after stretching.

The experimental data show that when the length of the membrane exceeds 10 μm, *η_eff_* rarely exceeds 5 pN•s/μm. In this case, tension decline rate was relatively fast, and the tension curve appeared distinctly different from that of large *η_eff_*. We found that the data after first two seconds of relaxation curves displayed a linear-like decline, which deviated from the model prediction. This deviation is similar to relaxation curves obtained from artificial liposomes and plasma membrane spheres, where people claimed the force relaxes with two timescales after elongation^32^.

The fast decline curve segment fitted in our model very well. Therefore, we corresponded them to the flow of phospholipids from membrane to tethers. We attributed the slow declining segment to the slow transportation of surrounding phospholipids to the region where the membrane tether is attached. The plasma membrane is separated into small regions at different levels by actin-based membrane skeleton, raft domain, and dynamic protein complex^20^. Cells use local invaginations and surface folds to form a membrane reservoir to buffer the rapid change of membrane tension during the stretching process^17,33^. Therefore, we believe the slow force decline indicates that membrane reservoirs have limited molecular interchanging with the surrounding membrane region. Once a great amount of lipid has been transported to the membrane tether, lipids diffuse to the region where the tether is attached. That will cause the slow decline on our relaxation curves.

In order to distinguish the fast decline process, we used data within the first two seconds of relaxation curves for fitting. The fitted *η_eff_* shows a clear plateau around 1.42 pN•s/μm, which we attributed to cases that no membrane protein stuck in the neck where membrane tethers attach. Since the curvature of the phospholipid membrane changes dramatically in the neck, we can reasonably suspect that protein molecules may be temporarily stuck in the border between membrane and tether until they got a chance to enter membrane tethers. Thus, big *η_eff_* might corresponded to cases that membrane proteins of different sizes and quantities stuck in the attachment point of the tether and blocked the free phospholipids flow. Relaxation curves divided into two parts with very different *η_eff_* might represents the process of stuck protein leaving the neck region and going into the tether. Therefore, we believed the measurement of *η_eff_* could help people understand the size and quantity of membrane proteins by fitting relaxation curves.

The relaxation curve method can obtain a large number of *f_0_* and *η_eff_* for statistical analysis. Using statistical analysis, we obtained the correlation between *η_eff_, f_0_* and tether length. We found that dynamic characteristics of long tethers significantly differ from that of shorter tethers. For example, big *η_eff_* presented more frequently on short tether than on tethers longer than 10μm. It indicates that the effect of membrane protein and membrane skeleton on relaxation curve might decrease as the tether length increase. For the same reason, small *f_0_* for long tether might be a signal of membrane reservoir depletion. Previous research only distinguish the tether formation stage and continuous extension stage^24,34^ On the contrary, our experiments show that the correlation between membrane length and physical parameters may contain detailed information of cell membrane dynamic changes.

Most of measured *η_eff_* in our experiments are around 1.42 pN•s/μm, which is about 10 times that of neuronal growth corn^23^. The force required to stretch out the filament on HeLa cells was about 130 pN, which is also much greater than the force (10-25 pN) normally required to stretch out the filament on neuronal growth corns. These phenomena indicate a great difference in surface tension and frictions among different cells and conditions. At this moment, fluorescent labels have been already employed in the measurement of tether diameter and protein distribution^13,35^. In the future, combining fluorescent labeling and relaxation curve method may help to observe the migration of proteins from cell membrane to tether, and understand the dynamics of cell membranes more comprehensively.

In summary, we characterized relaxation curves of HeLa cell membrane tether with established model. The relaxation curve method not only greatly improves the efficiency of optical tweezers in studying membrane tethers, but also enables the study of cytoskeleton and the dynamic process of membrane protein diffusion during tether elongation. The new approach to model relaxation curves provides a new perspective for studying the mechanical properties and dynamic process of cell membranes.

## Material and Methods

### Sample preparation and optical tweezers setup

Our optical tweezers setup^36^ focused a 1064 nm laser (Amonics, AFL-1064-40-R-CL) into a micro meter point through a 60x water immersion objective (Olympus UplanSApo, 60X, N.A. 1.2) to trap a two micro meter diameter polystyrene (PS) bead (Duke scientific Inc.). Laser light scattered by the bead was collected by a condenser and then projected onto a PSD (position sensitive device) in a back focal plane manner, so that the bead position signal was converted into electrical signal recognized by computer.

Firstly, 5 μl 2 μm diameter PS beads were washed with 500 μl PBS, and then dispersed in 50μl PBS. Next, 5 μl washed beads were added with 5 μl of 0.5 mg/ml lectin (lectin from Phaseolus vulgaris, Sigma L2646-25MG) and incubated at 4 °C for 1 hour. The chamber was made by inserting two separated narrow parafilm spacer between a glass slide and a cover glass. Then 30μl 10 μg/ml polylysine solution was injected into the chamber and left inside a moist box at room temperature for 30 minutes. After that, the chamber was washed with 400 μl PBS and 100 μl DMEM (Gibco) buffer in order. HeLa cells were in advanced DMEM (Gibco) with 10% fetal bovine serum (HyClone, Logan, UT), 100 units/ml penicillin (Gibco) and 100 μg/ml streptomycin (Gibco) at 37°C with 5% CO2. 5-10 μl cell culture medium was diluted with 50-100 μl DMEM (GIBCO) buffer and then injected into the sample chamber, and cultured at 4°C for 20 minutes. The lectin incubated PS beads were diluted with 100 μl DMEM (GIBCO) buffer, and then injected into the sample chamber right before experiment. The process was collected by CCD and displayed on the computer screen in real time.

### Model establishment

#### 1. Tension and radius of phospholipid tether

For a tube consisting of incompressible fluid, such as membrane tethers, the bending energy of the side wall is the major source of tension. Thus, the tension inside the tube correlates with the radius of tether curvature, which represents bending energy. In order to find the dependency of tube diameter and tension, we first considered a segment of phospholipid membrane tether with length L and radius *R_t_* under tension *f* (Fig, 5). The bending energy inside the segment is 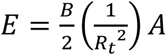, where *B* is the bending modulus of the membrane and *A* = 2*πR_t_L* is the surface area of the segment. If the membrane is incompressible, the surface area of membrane tether will be a constant when the diameter of tether changes. Then the change of free energy is

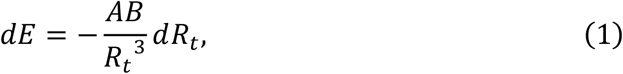

where *A* = 2*πR_t_L*, so that 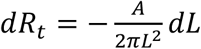, by introducing equation (1), gives

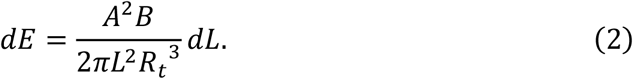

**Figure 5.**
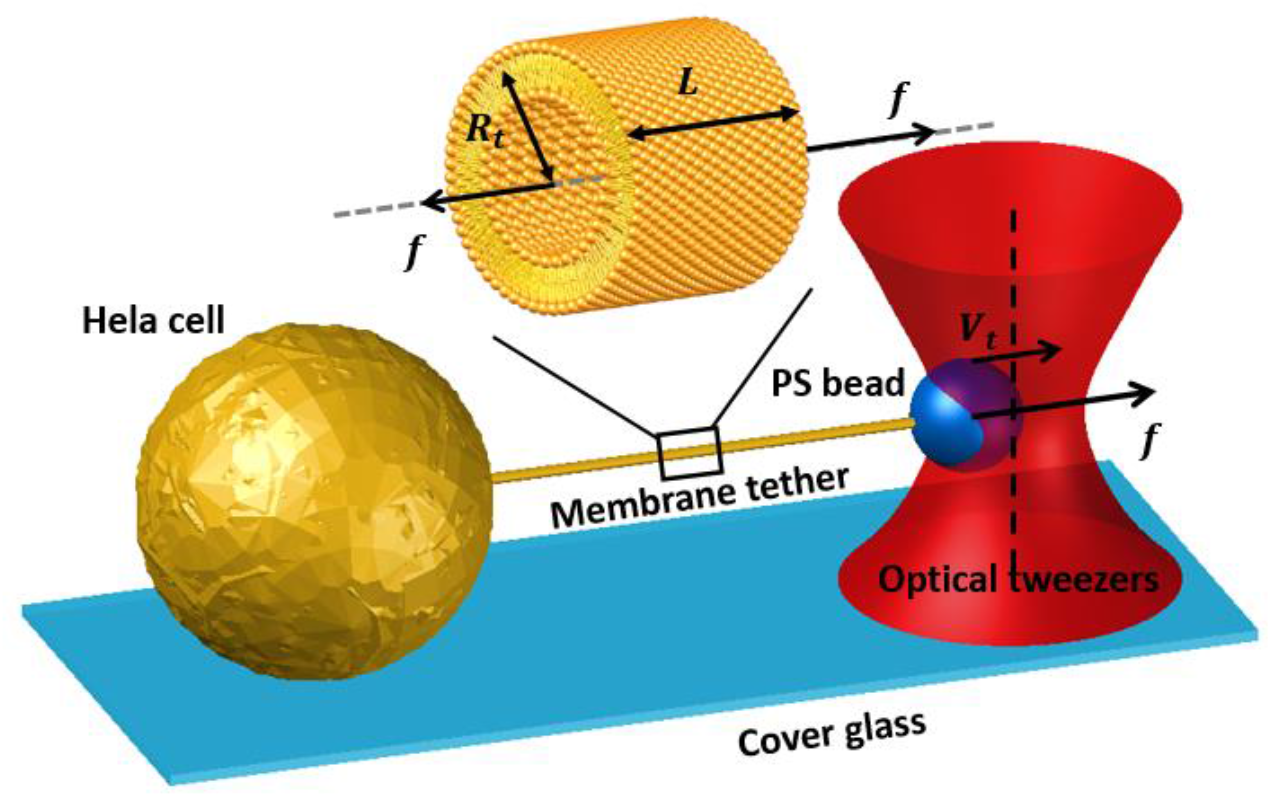
Schematic drawing of membrane tether relaxation. The big sphere is HeLa cell. The small sphere is PS bead captured by optical tweezers. The line between two spheres is membrane tether. The gray dash line indicates the central axis of membrane tether. *R_t_* is the radius of tether, *L* is the length of tether, *f* is the external force exerted by the optical tweezers on the bead and the tension on the membrane tether, and *V_t_* is the velocity of the bead.

According to the first and second laws of thermodynamics, the reversible isothermal work of a control volume *dW* = *fdL* equals to system free energy change dE for quasi-static processes. Combine it with equation (2) to eliminate *dL*, we get

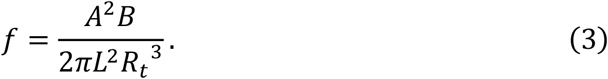

As *A* = 2*πRL*, equation (3) can be simplified to 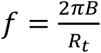, and the differential relationship between radius and force is obtained as follow:

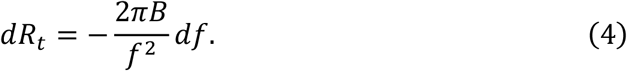

#### 2. Relaxation process of membrane tension under the control of optical tweezers

As showed in Figure 5, the overall force applied to the PS bead is balanced. Therefore, the pulling force from optical tweezers equal to tether tension *f*. It is well accepted that tension *f* equals to the sum of membrane tension, and the friction due to flowing phospholipids from membrane to elongating tether. When the active stretching stops, phospholipids flow tends to stop. Then the tension decreases due to declined friction. Thus, the tension is less than the optical tweezers pulling force and bead will be pulled toward the laser focus. Therefore, the membrane tether will keep elongating for a little while until the force applied on the bead balances with membrane tension and the phospholipids completely stops flowing (Fig. 1B). Therefore, the tension in the membrane tether exhibited a gradual decline curve from the dynamic tension *f* to the static tension *f_0_* when active stretching stops.

In order to evaluate the contribution of friction, we first consider phospholipids flow from the cell membrane to the membrane tether. At any point on the cell membrane, the velocities of the inner and outer phospholipids are expressed as

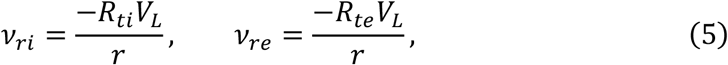

respectively, where *v_ri_* and *v_re_* are the flow velocities of inner and outer phospholipids at the radial position *r* on the cell membrane. *R_ti_* and *R_te_* are the radius of inner and outer membrane tether. *V_L_* is the velocity of lipid flow on the tether (Figure 6).

**Figure 3.**
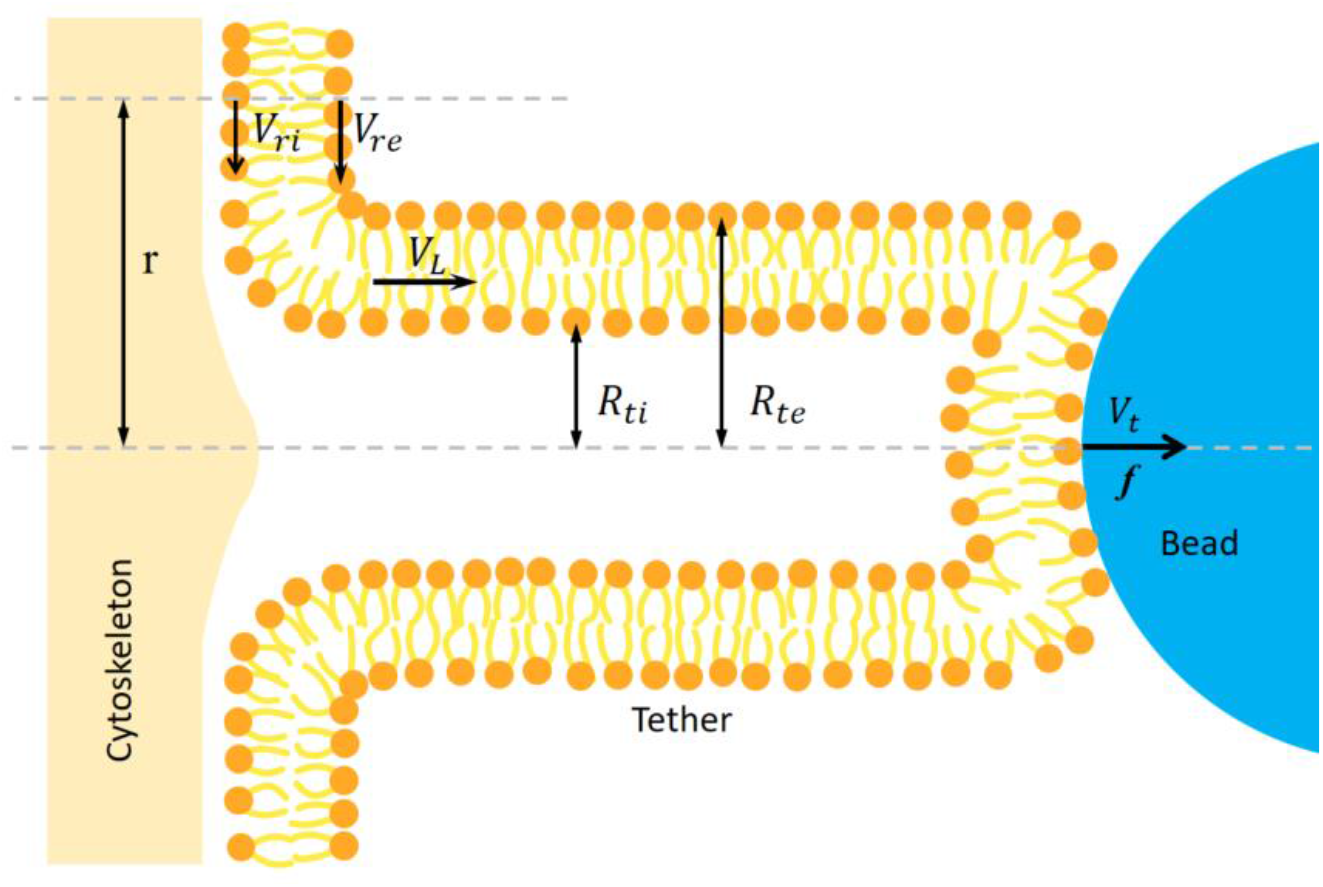
Schematic of an elongating tether. The shadow on the left side is cytoskeleton, phospholipid is pulling off a cytoskeleton, and the bead on the right side is pulled by optical tweezers under the force *f* The flow velocity of phospholipids on the cell membrane are *V_ri_, V_re_*, and the distance from the membrane tether central axis is *r. R_ti_* and *R_te_* are the radii of inner and outer tether. *V_L_* is the velocity of lipid flow on the tether.

When the rate of tether elongation *V_t_* does not change, the tension is a constant and *V_L_* equals to *V_t_*. However, when the tension in membrane tether is not a constant, the tether radius will change as described in equation (3). Therefore, extra lipid is needed to fill in and increase tether surface if the radius keeps increasing. Thus, the velocity of lipid flow can be written as *V_L_* = *V_t_* + *V_R_*, where *V_R_* represents the change of tether surface caused by radius increasing.

If the length of the membrane tether is *L*, and the radius is *R_t_*, we have 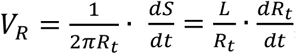, where *S* = 2*πR_t_* · *L* is the surface area of membrane tether. Put together we have:

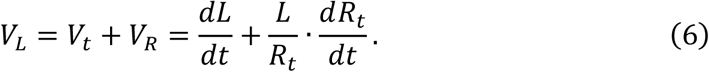

According to Hochmuch’s derivation^23^, the relationship between membrane tether tension *f* and phospholipid flow velocity *V_L_* can be written as follow:

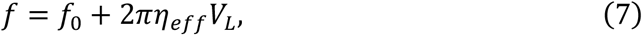

where *f_0_* is the static tension, *η_eff_* is effective viscosity coefficient, which represents the viscous resistance of phospholipids from cell membrane to membrane tether. The difference between equation (7) and equation (29a) in Hochmuch’s paper^23^ is that tether radius is a constant so that the velocity of lipid flow *V_L_* equal to the velocity of tether elongation *V_R_* in Hochmuch’s paper, but our *V_L_* includes an extra component *V_R_* associated with tether radius. Substitute equation (6) into equation (7) and we get

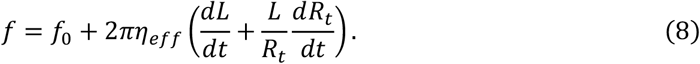

Then we look at the system of optical tweezers, bead, and membrane tether. Once active stretching stops, the distance between the cell and the optical tweezers is a constant. Thus, the tiny displacement of the bead to optical tweezers *dl* equals to the elongation of the tether *dL*. Therefore the elongation velocity is *V_t_* = *dL*/*dt* = *dl*/*dt*. For the optical tweezers, *dl* = −*df*/*k_Tw_*, where *k_Tw_* is the stiffness of optical tweezers. By substituting this relation and equation (4) into equation (8), we get the differential equation of tension *f* versus time t:

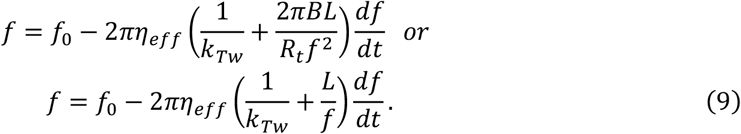

As the bead approaching the center of the optical tweezers, the force *f* exerted on the bead by optical tweezers and the velocity *V_t_* gradually decrease until reaching equilibrium. The maximum tension is recorded as *f_s_* at t=0; *f* equal to the static tension *f_0_* when *t* approaches to infinity. Substituting these boundary conditions into equation (9), the function of *f* and *t* is obtained by separating variables:

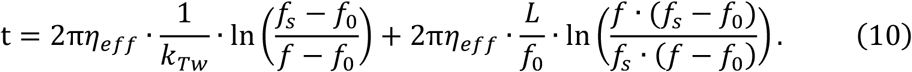

This is the relationship we use to fit relaxation curves of membrane tether tension and time, right after active stretching.

## Data availability

The data that support the findings of this study are available from the corresponding author upon reasonable request.

**Supplementary Information** is linked to the online version of the paper at www.nature.com/nature.

## Acknowledgments

This work was financially supported by the grants from the National Natural Science Foundation of China (grant 31870759, 61535011), the Fundamental Research Funds for the Central University (Grant number: WK9110000025), the National Cancer Center Climbing Funds (Grant Number:NCC201812B036), the Natural Science Foundation of Anhui Province (Grant Number:2008085MH288), the New Coronavirus Infection Emergency Science and Technology Project, Clinical Research Hospital of Chinese Academy of Sciences(Grant Number: YD9110002010).

## Author contributions

H. W. contributed to concept and design, experiment and manuscript writing; X. L. contributed to experiments and manuscript preparation; X. S. provided HeLa cells; M. L. contributed to manuscript discussion; Y. L. contributed to concept and manuscript writing.

## Author Information

Reprints and permissions information is available at www.nature.com/reprints.

### Competing interests

The authors declare no competing interests.

Correspondence and requests for materials should be addressed to.

