## Supplemental figure and table for "Characterizing mechanics of membrane tether from relaxation curve obtained by optical tweezers"

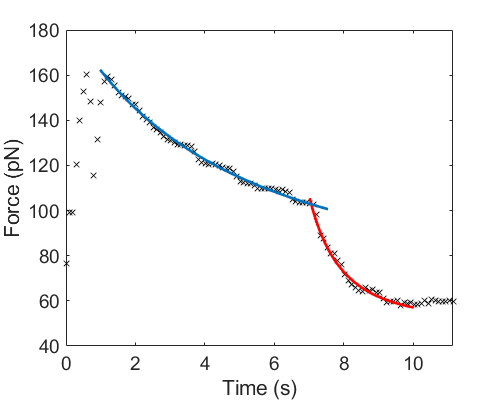

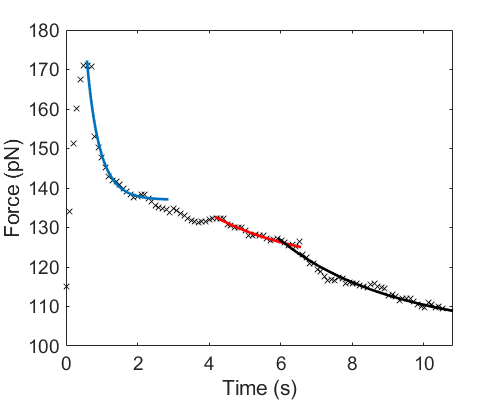

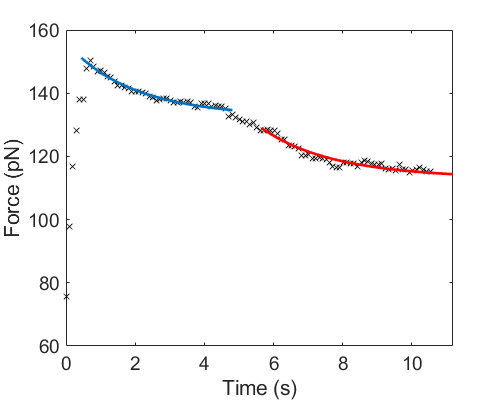

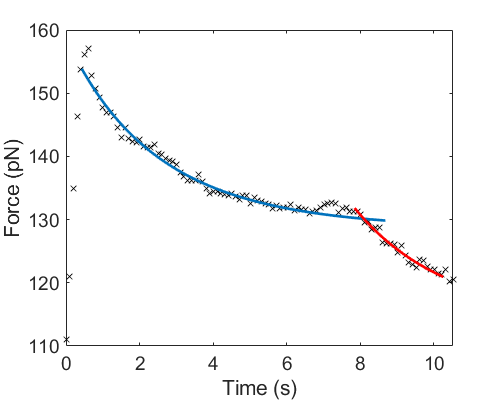

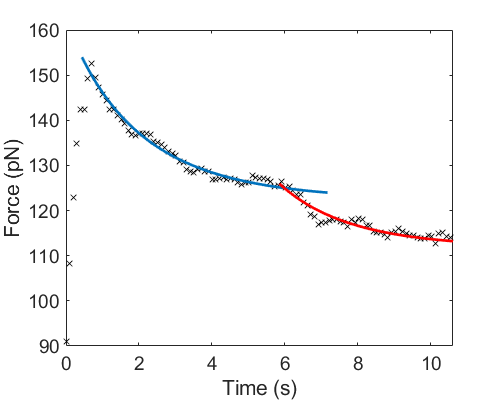

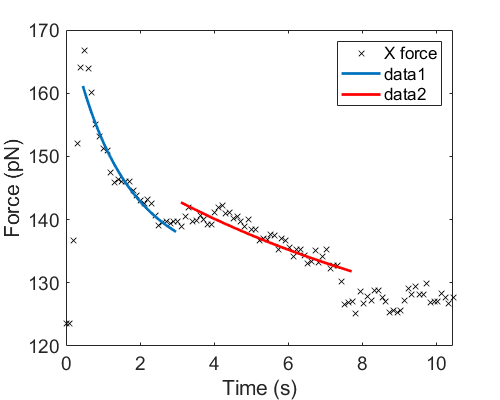

Figure S1. Relaxation curves with turning points. Black crosses are experimental data. Blue and red curves are fitting results.

Table S1. *η_eff_* and *f*_0_ obtained from fitting

| Segment number | Segment 1 | | Segment 2 | | Segment 3 | |
| --- | --- | --- | --- | --- | --- | --- |
|  | *η_eff_* (pN•s/μm) | *f*_0_ (pN) | *η_eff_* (pN•s/μm) | *f*_0_ (pN) | *η_eff_* (pN•s/μm) | *f*_0_ (pN) |
| 1520 | 51.7 | 70.0 | 6.5 | 52.4 |  |  |
| 1538 | 4.1 | 137 | 25.3 | 119.7 | 30.9 | 102.7 |
| 1541 | 12.4 | 132.4 | 11.7 | 113 |  |  |
| 1543 | 12.6 | 128.6 | 11.6 | 113.4 |  |  |
| 1547 | 7.0 | 122.4 | 5.8 | 111.9 |  |  |
| 1616 | 1.9 | 131.9 | 11.73 | 111.2 |  |  |
